# Early development of the glucocorticoid stress response in dyeing poison frog tadpoles

**DOI:** 10.1101/2024.05.31.596457

**Authors:** Lisa L. Surber-Cunningham, Lucas S. Jimenez, Lauren W. Mobo, Sarah E. Westrick, Eva K. Fischer

**Author notes:** Corresponding author: Lisa L. Surber-Cunningham, Department of Evolution, Ecology, and Behavior School of Integrative Biology, University of Illinois Urbana-Champaign 505 S Goodwin Ave, Urbana, IL 61801.

## Abstract

In vertebrates, the glucocorticoid “stress” response (corticosterone or cortisol) through the hypothalamic-pituitary-adrenal (HPA) axis influences many essential functions, including behavior, metabolism, immunity, and ontogenetic transitions. During development, stress responses can be adaptive if they facilitate antipredator behavior and modulate developmental speed to adjust to environmental conditions; however, these same responses can be maladaptive when energetic costs become too high and developmental speed trades-off with size and health at maturity. Thus, the timing of HPA-axis development may be aligned with specific developmental challenges and opportunities presented by a species’ life history strategy. In anurans (frogs and toads), corticosterone plays critical roles in development and behavior, and concentrations can fluctuate in response to environmental stressors. Given the role of corticosterone in ontogenetic changes and behaviors, we studied the development of the glucocorticoid stress response in tadpoles of the dyeing poison frog (*Dendrobates tinctorius*), a species with a unique life history that includes transport to water after hatching on land and aggressive and cannibalistic behavior. We measured the excretion rate and whole-body concentration of corticosterone and the corticosterone response to adrenocorticotropic hormone (ACTH) in free-swimming tadpoles after transport and throughout metamorphosis. We found no significant differences across development in excretion rates or whole-body concentration of corticosterone, nor corticosterone response to ACTH, indicating that that the glucocorticoid response develops early in ontogeny. This pattern differs from those in other species, suggesting the unique ecological pressures faced by *D. tinctorius* have shaped the development of the glucocorticoid stress response in this species. More broadly, this study illustrates how life history strategies and tradeoffs impact the timing of the development of stress responsivity.

## Introduction

Across development an essential challenge for organisms is to detect and respond to environmental challenges. In vertebrates, these responses are mediated in part via glucocorticoid “stress” hormones that regulate essential functions, including behavior, metabolism, immunity and ontogenetic transitions (Wada 2008, reviewed in Carr, 2011 and Landys et al., 2006). Given their wide-spread effects, glucocorticoids are also essential in coordinating phenotypic responses to stressors (Dantzer, 2023; Hau et al., 2016). Though the stress response is essential, chronically elevated glucocorticoids can cause problems including decreased immune function (Charmandari et al., 2005), elevated heart rate and blood pressure (Charmandari et al., 2005), negatively impacted growth rate (Glennemeier & Denver, 2002) and compromised cognitive ability later in life (Kitaysky et al., 2003). At an evolutionary timescale, the need to balance essential functions with negative effects leads to variation in when physiological stress responses develop (Wada, 2008).

Selection may mitigate tradeoffs between stress responsivity and hyperactivity by changing when the stress response develops (Wada, 2008). Many altricial species have minimal glucocorticoid reactivity early in life, a pattern thought to facilitate faster growth rates. For example, mice undergo a Stress Hypo-Responsive Period (SHRP) at four to fourteen days post birth, in which individuals have low basal stress hormone levels and an inability to mount a stress response to a mild stressor (Schmidt et al., 2002). However, in species with independent and active young, an early developed stress response may be crucial despite potential negative side effects. For example, precocial birds like turkeys show significant glucocorticoid activity from the day they hatch (Davis & Siopes, 1985), likely to facilitate immediate antipredator behaviors from their ground nests (Wada, 2008). In general, there is a balance between the timing and abundance of glucocorticoid production associated with life-history characteristics, developmental timing, and survival strategies across species.

In vertebrates, the neuroendocrine stress response is primarily mediated by the hypothalamic-pituitary adrenal (HPA) axis (Carr, 2011; Dantzer, 2023) or hypothalamic-pituitary-interrenal axis (HPI) in amphibians and fish (Denver, 2009a). When the HPA axis is activated, the hypothalamus releases corticotropin releasing hormone (CRH) that induces the pituitary gland (or anterior pituitary (Carr, 2011)) to secrete adrenocorticotropic hormone (ACTH). ACTH is transported to the adrenal cortex, which then releases glucocorticoid steroid hormones (Sapolsky et al., 1986). The glucocorticoid hormones cortisol and corticosterone are well-studied and – despite their many functions – are commonly referred to simply as ‘stress hormones.’ Though cortisol and corticosterone are generally considered functionally interchangeable, which glucocorticoid is dominant varies across species (Sapolsky et al., 2000).

The role of glucocorticoids in development is particularly interesting in the unique life-history of amphibians where they play a key role in metamorphosis (Denver, 1993, 1998, 2009b), and developmental trade-offs in response to environmental stressors are well-studied (McDiarmid & Altig, 1999). In many species of larval anurans (tadpoles of frogs and toads), corticosterone levels increase throughout development, peaking at metamorphosis (Jaudet & Hatey, 1984; Krug et al., 1983). These developmental shifts in corticosterone are in addition to fluctuations in response to a variety of well-studied environmental stressors that tadpoles face, including predation, competition, and habitat desiccation (Bókony et al., 2021; Denver, 1998; Forsburg et al., 2019a; Ledón-Rettig et al., 2023; Ledón-Rettig & Lagon, 2021). Acute changes in corticosterone impact tadpole antipredator behavior (Fraker et al., 2009; Kulkarni & Gramapurohit, 2017), morphology (Glennemeier & Denver, 2002; Hossie et al., 2010), and increase developmental speed (Denver, 1993, 1998, 2009b). Elevated corticosterone can increase developmental rate to rapidly escape a drying or dangerous pond (Pfennig et al., 1991), but rapid metamorphosis can have a negative impact on growth and survivorship later in life (Francesco Ficetola & De Bernardi, 2006). Thus, the unique life-histories and diverse environments of tadpoles highlight the potential for glucocorticoid mediated developmental tradeoffs.

We studied glucocorticoid activity and responsivity in tadpoles of the dyeing poison frog (*Dendrobates tinctorius*), a neotropical anuran species with several unique life history traits. Despite the variation in life histories, environmental pressures, and developmental rates across tadpole species, glucocorticoids and HPA responsivity have largely been studied in temperate, rapidly developing species. Compared to most temperature frog species that lay hundreds to thousands of eggs in the water and provide no care to eggs or offspring, adult *D. tinctorius* lay three to eight terrestrial eggs and provide parental care. Males clean, hydrate, and defend their eggs, and after hatching carry tadpoles “piggyback” to pools of water in fallen trees, trees with buttresses, and fallen palm bracts (Rojas, 2014; Rojas & Pašukonis, 2019). Once this transition from land to water is complete, tadpoles are independent from their parents and faced with predators, competition for resources, disease, habitat desiccation, and aggressive and cannibalistic conspecifics (Fischer et al., 2020; Rojas & Pašukonis, 2019). *D. tinctorius* tadpoles are able to survive in highly variable pool conditions (Fouilloux et al., 2021) and exhibit distinct behavioral responses to different environmental stressors from an early stage (Surber-Cunningham et al., 2024). Their developmental period is relatively slow, lasting a reported two months in the wild (Rojas, 2014; Rojas & Pašukonis, 2019) and three to four months the lab (personal observation).

Our study aimed to determine if glucocorticoid activity and responsivity shift during the ontogeny of *D. tinctorius* tadpoles, potentially facilitating developmental transitions and supporting key physiological and behavioral adaptations. Given the shared and distinct life history features between *D. tinctorius* tadpoles and more temperate species, we had two alternative predictions. On the one hand, given the constraints of metamorphosis and environmental challenges shared with other tadpoles (predation, desiccation, etc.), *D. tinctorius* tadpoles’ stress response could have a developmental trajectory similar to other species, with a peak in corticosterone close to metamorphosis. Alternatively, we predicted *D. tinctorius* might have an early-developed glucocorticoid stress response to facilitate the behavioral dexterity involved in coordinated transport, the ontogenetic transition from land to water, responses to environmental challenges immediately upon arrival in pools, and an extended developmental period. To test these alternatives, we compared (1) the excretion rate and whole-body concentration of corticosterone and (2) the corticosterone response to ACTH across developmental stages. We additionally confirmed corticosterone as the dominant glucocorticoid and validated waterborne hormone sampling methods previously used in adult *D. tinctorius* (Rodríguez López, 2015; Westrick et al., 2023) in tadpoles.

## Methods

### Animal husbandry

*D. tinctorius* tadpoles used in this study were hatched in the Fischer Lab frog colony at the University of Illinois Urbana-Champaign (UIUC). Once parents deposited a tadpole into a pool, we moved the tadpole to a transparent plastic cup (6cm diameter, 7.5cm height) with mesh bottom within a larger plexiglass aquarium. We separated tadpoles into three aquaria with tadpole densities of 12 – 20 individuals per aquarium in approximately 20 liters of reverse osmosis (RO) water. In each aquarium, tadpoles shared water through the mesh bottoms of each plastic cup but were unable to physically interact. We kept the water temperature constant at 22.8 ±1°C and kept the room under a 12:12-hour light/dark cycle. Tadpoles were also given 15 minutes of UV light per day to support development (Pahkala et al., 2003). We fed tadpoles a diet of shrimp flakes and rabbit food pellets three times per week. We performed daily health checks, weekly partial water changes, and weekly water quality checks (pH, salinity, temperature). All animal care and experimental procedures were approved by the Animal Care and Use Committee of the University of Illinois at Champaign-Urbana (IACUC Protocol number 20147).

### Staging & morphological measurements

We measured tadpoles’ total body mass, body length, tail length, head width, and Gosner stage. To measure wet mass, we tared a portable scale (Weightman model B200) with a small cup of water, and a tadpole was gently placed into the cup via a dry spoon. We recorded mass in grams to the nearest 0.01 g. To measure total body length, tail length, and head width, we placed tadpoles on a millimeter grid, and straightened body position gently with a fine paint brush.

Because tadpole developmental speed is highly plastic (Pfennig et al., 1991), it is standard practice to use developmental stage rather than age. We categorized tadpole development via Gosner stage, a numeric system based on key morphological characteristics (Gosner, 1960). We examined tadpoles under a stereo microscope (Leica S9E) for accurate Gosner staging. We separated tadpoles into three developmental categories: early stage (no external limb development, Gosner stages 25-29, n=10), middle stage (subtle external leg development, Gosner stages 30-34, n=10), and late stage (prominent external leg development, Gosner stages 35-40, n=10). Tadpoles were out of the water for no longer than five minutes during morphometric data collection, and no tadpoles were harmed in the process.

### ACTH hormone challenge

To test for differences in the physiological stress response, we manipulated the HPA axis via an ACTH challenge. ACTH challenges are used across vertebrate taxa to stimulate the HPA axis and assess stress responsivity (Hammond et al., 2019; McClelland & Woodley, 2021; Murray et al., 2020). ACTH injections took place one or two days after morphological measurements. We utilized a within-individual repeated measures design in which tadpoles were injected with both an intraperitoneal injection of 0.5 ug ACTH per gram body weight (Sigma-Aldrich®, product no. A7075) (Forsburg et al., 2019) diluted in 0.9% saline and a control injection of 0.9% saline (with volume matched to the ACTH injection), with six days between injections. We used a 50µL Hamilton syringe (Gastight® #1705) with single use 30G hypodermic needle (Excel Int, model 30Gx1) for injections. During injection, we placed tadpoles on a clean wet sponge and held them in place with a wet strip of paper towel.

### Waterborne hormone collection

We used waterborne hormone collection as a minimally invasive method for repeated hormone sampling in small-bodied amphibians. To collect each hormone sample, we soaked the tadpole in 50mL RO water in a ten-centimeter diameter cylindrical glass container for 1 hour. To measure pre-injection glucocorticoids, we collected a sample immediately prior to injection. To measure how glucocorticoids respond to an ACTH and control injections over time, we collected water samples at four time points post-injection: one hour, two hours, four hours, and twenty-four hours. Pre-injection hormone collection started between 9am and10am, with injections one hour after (between 10am and 11am). All individuals completed an ACTH trial and a saline trial one week apart; thus, we collected ten waterborne hormone samples per individual (five per trial).

The order of the injections (ACTH/control) was counter-balanced to control for order effects. After collection, we filtered water samples through filter paper (Whatman 1001-125) to remove any debris and stored the filtered samples in 50mL conical tubes at -20°C until steroid hormone extraction (see below). We collected a total of n=295 waterborne hormone samples (n = 30 tadpoles). One individual died between week one and two, but death believed unrelated to experiment and data points still included.

### Whole body hormone extraction

We euthanized tadpoles immediately following the 24-hour hormone collection using an overdose of sodium bicarbonate buffered tricaine methane sulfonate (MS-222), a commonly-used anesthetic for fish and amphibians (Ramlochansingh et al., 2014; Topic Popovic et al., 2012). We rinsed bodies thoroughly with RO water, and stored bodies individually in microcentrifuge tubes at -80°C for two weeks. Prior to whole-body hormone extraction, we weighed bodies to the nearest 0.001 g using a microbalance (Mettler Toledo ML203T/00).

Whole bodies were homogenized in liquid nitrogen using a mortar and pestle. We transferred homogenized tissue to a pre-weighed 15 mL centrifuge tube and recorded the powder-only mass. Following weighing, we added 5 mL of methanol and vortexed for about 30 seconds. Samples were stored overnight at -20 °C. The next day we centrifuged tubes with methanol and tissue (4°C, 2000 rpm) for fifteen minutes (Eppendorf centrifuge 5810R). We carefully pipetted the supernatant (∼ 5 mL) into a 50 mL conical tube and added 45 mL of RO water. Samples were then processed through the solid phase extraction process via the vacuum manifold procedure (see below). We collected N=29 body samples for hormonal analysis (early=10, middle=9, late=10).

### Steroid hormone extraction

We extracted hormones from water samples and methanol-extracted whole-body samples using solid-phase extraction via C18 cartridges (Waters Corporation) on a vacuum manifold. We primed the cartridges with 6 mL of 100% methanol, rinsed with 6 mL RO water, and then passed the 50mL sample through. We used 4 mL of 100% methanol to elute the collected total steroid hormones into 13×100 mm borosilicate glass vials. We dried samples on a 37°C heating block under a stream of nitrogen gas. After completely dry, we covered tubes with tight fitting caps and stored at -20°C until quantification.

### Enzyme linked immunosorbent assay

Methods for quantification of waterborne hormone samples using enzyme immunoassay (EIA) have been utilized and validated in various amphibians (Baugh et al., 2018; Forsburg et al., 2019; Gabor et al., 2016). These methods have been successfully applied in *D. tinctorius* by our research group (Westrick et al., 2023) and others (Rodríguez López, 2015). We previously confirmed parallelism between standards provided by the manufacturer and water samples from *D. tinctorius* for corticosterone and cortisol (Westrick et al., 2023).

To quantify the concentration of cortisol and corticosterone in each sample, we used DetectX® kits from Arbor Assays™. We reconstituted samples by adding assay buffer (catalog no. X065) to the dried waterborne (500 μL) and whole-body samples (750 μL) in borosilicate glass vials. We ran each sample in duplicate on both a corticosterone and a cortisol EIA. Individuals that completed all sample collections had a total of ten waterborne samples and one whole-body sample. Because we were interested in within-individual differences, we set up plates so that all the samples taken from an individual were run on the assay to reduce potential variation caused by plate-to-plate variation. We followed manufacturer guidelines for running all EIA plates. We prepared the standard curve and samples using the high sensitivity 100 µL assay format, such that the lowest point of the curve measured 19.531 pg/mL. The lowest point of the cortisol curve measured 50 pg/mL. We measured the optical density at 450 nm for all assays. The manufacturer reported cross reactivity of the cortisol EIA with corticosterone as 1.2%, and cross reactivity of the corticosterone EIA with cortisol as 0.38%.

To generate a standard curve for each plate, we used MyAssay software (www.MyAssay.com) to fit a four-parameter logistic curve (4PLC) regression. We used this software to calculate the concentration of glucocorticoids (pg/mL of assay buffer) for each sample based on the standard curve and averaged across duplicates. To calculate the excretion rate for water samples, we multiplied the concentration by the volume of assay buffer used to reconstitute the sample (0.5 mL) to get mass (pg) of the hormone in the sample. We then divided by the collection time (one hour) to get the release rate (pg of hormone per hour of collection). For whole-body samples concentrations, we corrected for the milliliters of assay buffer used (0.75 mL) and the mass of powder collected after grinding (pg of hormone per gram of dry body mass).

We removed samples with a coefficient of variation (CV) greater than 20% (n = 17 out of 328 total corticosterone measurements, n=233 out of 328 total cortisol measurements). Most cortisol samples fell at the low end of the plate measurement range, and high CVs for cortisol arose in part from increased noise among duplicates at the extremes of the detectable range. The remaining average intra-assay CV across all samples was 6.85% for corticosterone and 10.01% for cortisol.

To test the extraction efficiency of our hormone analysis, we collected additional samples (N=12) by soaking tadpoles in RO water for one hour. We divided each sample equally to measure the concentration of a control and a spike. For the spiked samples, we added either a cortisol standard (N=6) or a corticosterone standard (N=6). These samples were then processed through the extraction, drying, and plating methods described previously. To determine the “observed” concentration of the spiked standard, we subtracted the control sample’s value from the corresponding spiked sample’s value. The recovery rate was calculated by dividing the observed concentration by the expected concentration (i.e., the amount added to the spike) and multiplying by 100. Our results showed average recovery rates of 108% (range: 96.7%–127%) for cortisol and 87.5% (range: 78.5%–102%) for corticosterone.

### Statistical analysis

All statistical analyses were performed with R (v. 2022.07.1 + 554 R Core Development Team, 2022) in Rstudio (v. 2022.12.0 Rstudio Team, 2022). We log transformed corticosterone and cortisol release rates for all tests to achieve normality. For all analyses described below, we ran linear mixed models using the lmer() function in ‘lme4’ package (Bates et al., 2015), and calculated test stats and p-values using the package ‘lmertest’(Kuznetsova et al., 2017). To analyze if there were significant inter-plate differences, we ran separate models for corticosterone and cortisol with the glucocorticoid concentration as the dependent variable, plate number as a fixed effect and individual as a random effect. The variation between EIA plates was non-significant for corticosterone (F_1,30.9_=0.367, p=0.716) and cortisol (F_1,17.08_=0.017, p=0.898).

To analyze which glucocorticoid (cortisol/corticosterone) was released at a higher rate, we ran linear mixed effects models with the pre-injection glucocorticoid release rate (pg/hr) as the dependent variable, and glucocorticoid type (cortisol/corticosterone), developmental stage (early/middle/late), and the interaction between glucocorticoid type and developmental stage as fixed effects, and individual as a random effect. We ran linear models using the whole-body hormone concentrations to understand the abundance within the body, with concentration of glucocorticoids (pg/g) as the dependent variable, and glucocorticoid type, treatment (ACTH/saline control), developmental stage, and all interactions as fixed effects.

To assess pre-injection differences in corticosterone across developmental stages, we ran linear mixed effects models with the pre-injection glucocorticoid release rate (pg/hr) as the dependent variable, and developmental stage (early/middle/late) as a fixed effect, and individual as a random effect. Tadpoles underwent two trials (one ACTH and one saline injection) one week apart, thus we originally included week in this model, but removed it when nonsignificant (F_1, 26.1_=0.218, p=0.645). We ran a linear model using the whole-body hormone concentrations to understand the abundance within the body, with concentration of glucocorticoids (pg/g) as the dependent variable, and developmental stage as a fixed effect. We did not include body-size as a covariate in our models as it was confounded with developmental stage (later stage tadpoles are larger) and created problems with collinearity in the models.

We analyzed if whole-body corticosterone concentration predicted waterborne corticosterone release rate via linear model, with week two 24-hour waterborne release rate (the sample time point immediately preceding the body collection) as the dependent variable and the whole-body corticosterone concentration, developmental stage (early/middle/late), and their interaction as fixed effects. We did not include treatment in this model, because there were no differences between treatments at this time point.

To analyze if corticosterone increased in response to ACTH and if this response changed across developmental stages, we ran a linear mixed effects models with corticosterone concentration as the dependent variable. We included all water-borne sampling time points and fit models with treatment, timepoint, developmental stage, and all interactions between as fixed effects and individual ID as a random effect. We included sampling week as a fixed effect in original models but removed it to prevent over-fitting because it was not statistically significant (F_1, 238.26_=0.074, p=0.786). We ran a post-hoc analyses using the package ‘emmeans’ (Lenth et al., 2022) with pairwise tests to compare across timepoints, to compare the timepoints within each treatment group (timepoint x treatment), to compare the treatment groups within timepoint (timepoint x treatment), and the developmental groups between treatments (treatment x development). We did not run analyses testing the response of cortisol to ACTH because small sample sizes resulting from low cortisol values (see above) made robust statistical analysis impossible (N<=3 for 19/30 experimental groups of treatment*time*developmental stage).

## Results

Corticosterone release rate was significantly higher than cortisol release rate pre-injection (F_1,59.201_=189.475, p<0.0001, Figure 1A) and in whole-bodies (F_1,34=_46.546, p<0.0001, Figure 1B) in all developmental stages. Given the approximately seven-fold greater abundance of corticosterone and the high number of cortisol samples dropped due to low concentration (see Methods), we report differences across development and in response to ACTH challenge for corticosterone only.

**Figure 1.**
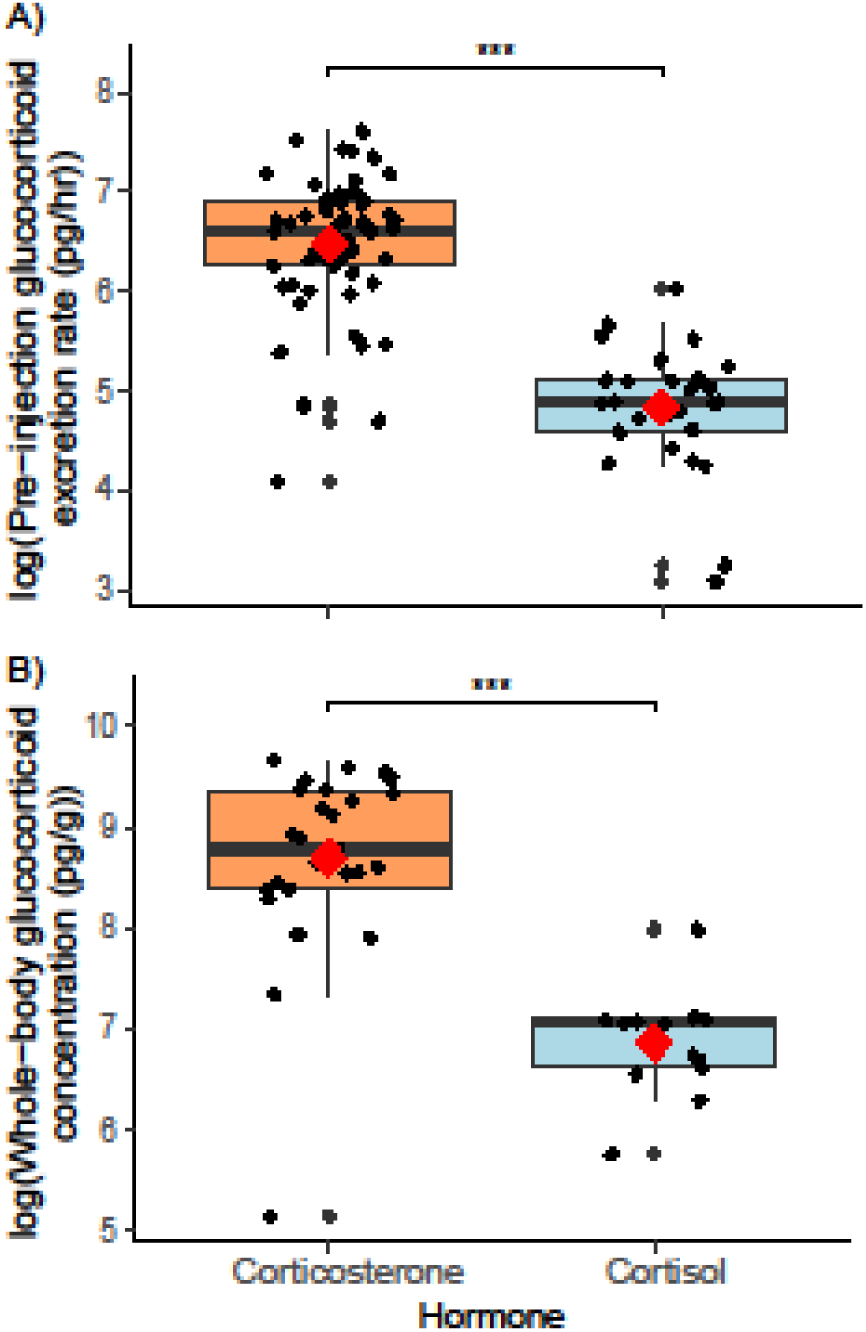
Tadpoles excrete more corticosterone (orange) than cortisol (blue) (A) pre-injection (B) whole-body. We collapsed developmental groups together because there were no differences between them. Boxplots show the median, the first and third quartiles, and whiskers as 1.5x the interquartile range. Black dots show individual data points. Red diamonds show group means.

Corticosterone release rate did not differ across developmental stages in pre-injection waterborne samples (F_2,47.04_=0.361 p=0.699) (Figure 2A) or whole-body samples (F_2,25_=1.043, p=0.370) (Figure 2B). Whole-body corticosterone predicted waterborne corticosterone (F_1,18_=21.95, p<0.005), and this relationship did not differ across developmental stages (F_2,18_=0.506, p=0.611) (Figure 2C).

**Figure 2.**
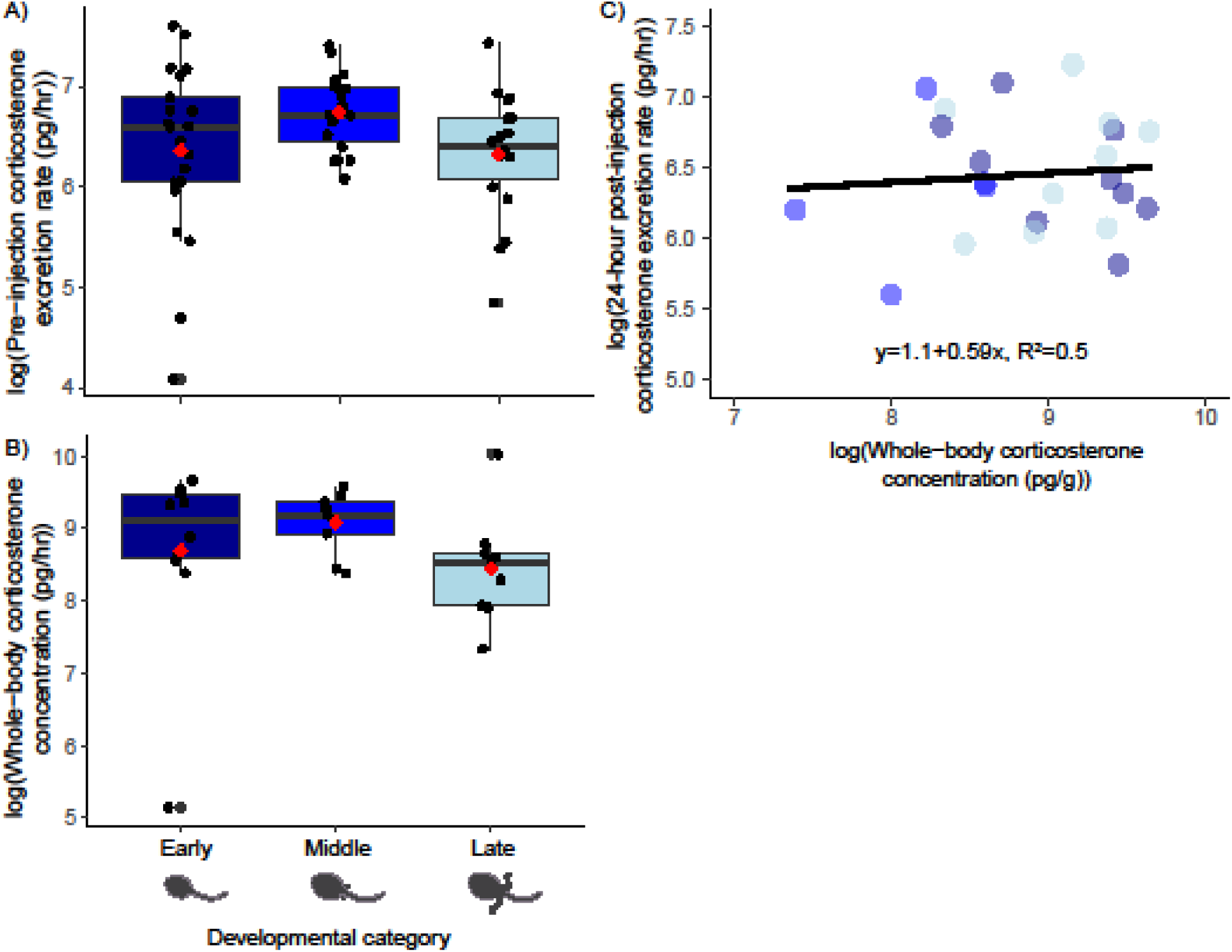
Early (dark blue), middle (blue), and late-stage (light blue) tadpoles do not differ in corticosterone in (A) pre-injection excretion rate or (B) whole-body concentration. Boxplots show the median, the first and third quartiles, and whiskers as 1.5x the interquartile range. Black dots show individual data points. Red diamonds show group means. (C) Whole-body corticosterone predicted corticosterone excreted, with no significant differences in the water-body relationship between developmental groups.

When we compared corticosterone responses in ACTH and saline treatment groups, we found an overall higher corticosterone excretion rate in the ACTH treatment group (treatment: F_1,230.6_=4.45, p<0.05) (Figure 3A). Sampling time point also had a significant effect on corticosterone excretion rate (time: F_4,224.94_ = 24.85, p <0.0001), though the pattern of change was the same in both treatment groups (treatment*time: F_4,224.84_ = 1.43, p=0.2243) (Table 1).

**Figure 3.**
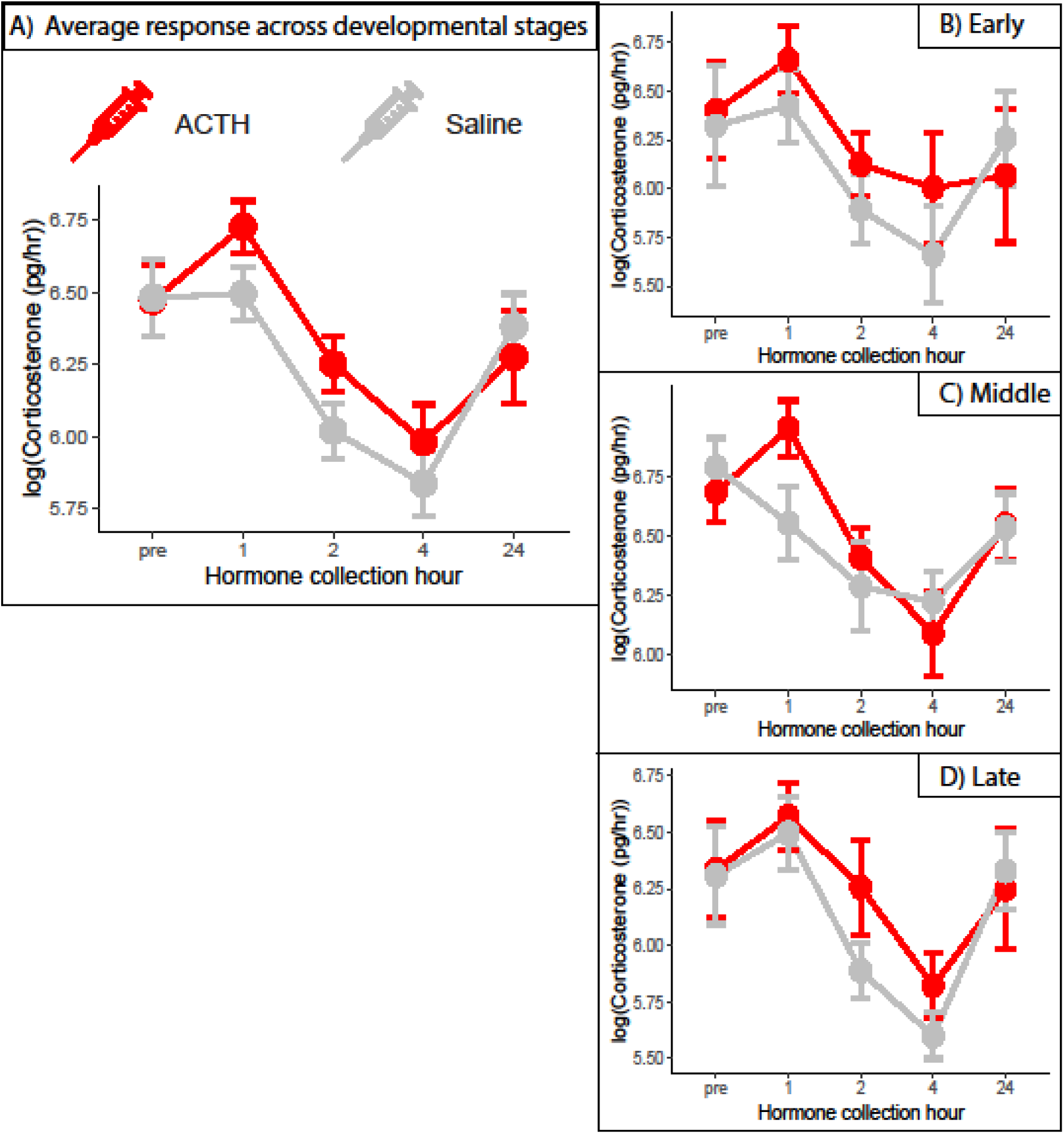
When (A) averaged across all developmental groups, tadpoles injected with ACTH (red) excreted significantly more corticosterone one hour and marginally more corticosterone two hours post injection compared to the saline control (gray). There was no significant difference in ACTH response between (B) early, (C) middle, or (D) late-stage tadpoles. Data are shown as means +/- standard error.

**Table 1.** Post-hoc results comparing corticosterone excretion rates pre and post injection samples, separated for ACTH and saline treatments.

|  |  |  |  |  |
| --- | --- | --- | --- | --- |
| Collection hours | ACTH |  | Saline |  |
|  | t | p | t | p |
| Pre - 1 | t<sub>225</sub>= -2.362 | 0.1297 | t<sub>224</sub>=-0.169 | 0.998 |
| Pre - 2 | t<sub>225</sub>= 1.926 | 0.3065 | **t<sub>225</sub>= 3.990** | **<0.001** |
| Pre - 4 | **t<sub>225</sub>= 4.367** | **<0.0005** | **t<sub>224</sub>= 5.848** | **<0.0001** |
| Pre - 24 | t<sub>225</sub>= 1.546 | 0.5337 | t<sub>225</sub>=1.033 | 0.8396 |
| 1 - 2 | **t<sub>225</sub>= 4.245** | **<0.0005** | **t<sub>225</sub>= 4.157** | **<0.0005** |
| 1 - 4 | **t<sub>225</sub>= 6.686** | **<0.0001** | **t<sub>224</sub>= 6.016** | **<0.0001** |
| 1 - 24 | **t<sub>225</sub>= 3.874** | **<0.005** | t<sub>225</sub>=1.197 | 0.7530 |
| 2 - 4 | t<sub>225</sub>= 2.369 | 0.1276 | t<sub>225</sub>=1.788 | 0.3833 |
| 2 - 24 | t<sub>225</sub>= -0.394 | 0.9949 | **t<sub>225</sub>= -2.861** | **<0.05** |
| 4 - 24 | **t<sub>224</sub>= -2.795** | **<0.05** | **t<sub>225</sub>= -4.653** | **<0.0005** |

Comparing between treatment groups, corticosterone release rate was higher in ACTH treated than saline treated tadpoles at one hour post injection (t_,226_ = 2.138, p<0.05) and was marginally higher at two hours post injection (t_1,227_ = 1.932, p=0.055) (Table 2). We found no effect of developmental stage on treatment response (treatment*stage: F_4,231.64_ = 0.20, p=0.8167), corticosterone changes across time (time*stage: F_4,224.93_ = 0.36, p=0.9389; treatment*time*stage: F_4,224.84_ = 0.76, p=0.6393), or their interaction (treatment*time*development: F_8,224.84_=0.759, p=0.639) (Figure 3B, 3C, 3D).

**Table 2.** Post hoc comparisons of corticosterone excretion of ACTH and saline across the hormone time collections.

|  |  |  |
| --- | --- | --- |
| Collection hour | corticosterone |  |
|  | t value | p value |
| Pre-Injection | T<sub>226</sub> = -0.050 | 0.9600 |
| Post 1 hour | **T<sub>1,226</sub> = 2.138** | **0.0336** |
| Post 2 hour | **T<sub>1,227</sub> = 1.932** | **0.0546** |
| Post 4 hour | T<sub>1,226</sub> = 1.335 | 0.1832 |
| Post 24 hour | T<sub>1,226</sub> = -0.534 | 0.5937 |

## Discussion

A major way vertebrates coordinate responses to environmental stressors is through the glucocorticoid stress response mediated by the hypothalamic-pituitary adrenal axis (Carr, 2011; Dantzer, 2023, Denver, 2009a). However, what constitutes an appropriate stress response can change across development as morphological and energetic changes shift the potential tradeoffs imposed by excessive or insufficient glucocorticoid activity. This shifting landscape may in turn shape evolved differences in the developmental timing of glucocorticoid stress responses specifically and HPA axis activity generally based on a species’ life history. In this study we investigated glucocorticoid activity and responsivity in dyeing poison frog (*Dendrobates tinctorius*) tadpoles, a species with a unique life history that includes transport to water after hatching on land, as well as aggressive and cannibalistic behavior. We specifically tested the alternatives that *D. tinctorius* would (1) exhibit a developmental pattern similar to most tadpoles, with increased glucocorticoid abundance just before metamorphosis (Glennemeier & Denver, 2002; Jaudet & Hatey, 1984; Krug et al., 1983), or (2) have an early-developed glucocorticoid stress response given the importance of glucocorticoids in ontogenetic transitions (Wada, 2008) and for rapid phenotypic responses to short-term environmental stressors (Dantzer, 2023; Hau et al., 2016).

We found no differences across tadpole developmental stages in corticosterone excretion rate in waterborne samples, whole-body concentration of corticosterone, nor ACTH response, supporting the prediction that *D. tinctorius* tadpoles have an early developed and consistently active glucocorticoid stress response. This result contrasts with findings in other species, where late-stage tadpoles have corticosterone levels two to three times higher than early or middle-stage tadpoles (Glennemeier & Denver, 2002; Jaudet & Hatey, 1984; Krug et al., 1983). Notably, other species that have been studied tend to be explosive breeders that produce 10s to 1000s of aquatic eggs and provide no parental care. We suggest these key life history differences could contribute to our findings here.

Our study also adds to the mounting evidence that there is species variation in glucocorticoid and HPA axis reactivity across development in tadpoles. In American bullfrogs (*R. catesbeiana*), late-stage tadpoles mount a response greater than early stage tadpoles (Krug et al., 1983), while Northern leopard frogs (*Rana pipiens*) and African clawed frogs (*Xenopus laevis*) have a larger ACTH response as early-stage tadpoles than late-stage (Glennemeier, 2002). In *D. tinctoirus*, we found equal reactiveness to ACTH across development. Below we discuss multiple, non-mutually exclusive reasons why early-developed and consistently responsive glucocorticoid activity could be important in this species.

We posit that the early development of the glucocorticoid stress response in *D. tinctorius* tadpoles may facilitate transportation and transition from terrestrial hatching sites to developmental pool, and once there, enable plastic responses to highly variable environmental conditions and conflicts with conspecifics. In *D. tinctorius*, tadpoles hatch terrestrially, then crawl onto the backs of their parents and hold on as they are transported for minutes to hours to their new aquatic habitat. This action takes collaboration between parents and offspring, as well as water balance while in the open air and during transition to a fully aquatic environment. An early developed stress response may facilitate these metabolically demanding behaviors (Magomedova & Cummins, 2016). We did not compare glucocorticoid abundance or responsivity pre-transport in this study, as out non-invasive waterborne sampling method would have inherently confounded measuring the effects of the terrestrial to aquatic transition. However, this is an exciting avenue for future research as this land-to-water transition is fundamentally different from the majority of frogs that lay aquatic eggs and do not provide care.

After arriving at their pond, *D. tinctorius* face challenges similar to those of most tadpoles (predation, competition, desiccation) as well as some additional challenges. The pools *D. tinctorius* tadpoles inhabit vary considerable in water quality, predator and conspecific abundance, and pool stability, especially compared to other neotropical species (Fouilloux et al., 2021). These variables are known to cause plastic developmental shifts in other tadpoles (McDiarmid & Altig, 1999), and such plasticity could be facilitated by an early developed glucocorticoid stress response and HPA axis in *D. tinctorius*. Additionally, *D. tinctorius* tadpoles are aggressive and cannibalistic (Fouilloux et al., 2022; Fischer et al., 2020), which makes interactions with conspecifics a potential danger or a potential meal. Indeed, we previously demonstrated that *D. tinctorius* exhibit behavioral responses to injured conspecific cues that are distinct from responses to food or predator cues (Surber-Cunningham et al., 2024). We suggest that corticosterone may be mediating behavioral responses to these unique cues as glucocorticoids are known mediators of behavioral plasticity generally (Dantzer, 2023), and are positively correlated with aggression (Summers & Winberg, 2006) and antipredator behavior (Fraker et al., 2009) in other tadpole species. Our findings lay the foundation for future studies to investigate the role of corticosterone as a mediator of phenotypic plasticity and juvenile aggression.

In general, tadpole responses to ACTH challenge were as expected: treatment groups did not differ pre-injection, corticosterone excretion was relatively higher following ACTH treatment, and corticosterone levels recovered to pre-injections concentrations 24 hours post-injection. While we expected this general response, we further asked whether the shape or magnitude of the response differed across development. We found no significant differences in ACTH response across developmental stages. This observation has two important implications.

First, it demonstrates that *D. tinctorius* tadpoles can mount a physiological stress response even early in development, confirming our prediction that this species has an early developed HPA axis. Second, these results indicate that developmental effects of corticosterone do not interfere with stress responsivity in this species. Taken together, these observations demonstrate that *D. tinctorius* can respond rapidly and consistently to stressors throughout their aquatic developmental period.

Although the response to ACTH was as we expected, one unanticipated result in our study was that corticosterone excretion pre-injection did not differ significantly from excretion one-hour post-injection in either the ACTH or saline group. This suggests both groups exhibited some handling stress associated with waterborne hormone collection (moving from their home cup to the hormone collection cup), as has been observed in other species (Wong et al., 2008).

Though waterborne hormone sampling requires some handling (Wong et al., 2008) and is inherently an integrative measure across time, we expect any effects of sampling procedure to be consistent across experimental groups. Similarly, any stress resulting from injection is shared in ACTH and control treatments. Importantly, we nonetheless found that corticosterone was elevated in the ACTH group as compared to saline, indicating that the HPA axis was activated by our pharmacological manipulation above responses to handling and/or injection. Indeed, when treated with ACTH, individuals had elevated corticosterone and corticosterone decreased more slowly than when treated with saline. At present, we cannot distinguish whether the lower corticosterone at 2-hours and 4-hours post injection results from short-term habituation to sample collection or negative feedback from the HPA-axis, and either alternative suggests that all tadpoles experienced at least a slight activation in the sampling and injection process. We note that we have no indication for habituation at longer timescales, as corticosterone levels at 24 hours post injection and in the second week of sampling were similar to those pre-injection and in the first week. While these observations underscore the importance of considering habituation procedures when conducting experiments involving glucocorticoid manipulation in tadpoles and other organisms, they do not negate the results and interpretations discussed here.

We found that whole-body corticosterone concentration predicted the release rate of waterborne corticosterone, and that this relationship did not change across development. From a technical perspective, our findings validate the use of waterborne hormone measurements in tadpoles of *D. tinctorius*. Waterborne hormone sampling is minimally invasive, allowing for repeated sampling in small-bodied individuals and non-destructive sampling of wild populations, thereby providing many potential avenues for future study. While correlations between body and water samples are expected, confirmation of this pattern is also biologically interesting, as previous studies have documented changes in body/water correlations across tadpole development (McClelland & Woodley, 2021). McClelland & Woodley (2021) report a lack of correlation between waterborne and plasma hormone concentrations in early-stage tadpoles and those undergoing metamorphic climax, suggesting excretion rate can be plastic across development, at least in some species. Data from additional species are needed to understand whether and how these patterns may be related to developmental changes in glucocorticoid responsivity and HPA axis activity.

Finally, our study confirmed that corticosterone is the dominant and most abundant glucocorticoid in *D. tinctorius*. Across tadpole development, corticosterone was approximately seven times more abundant than cortisol. Corticosterone is generally considered the primary glucocorticoid in amphibians (Denver, 2009a; Glennemeier & Denver, 2002; Hossie et al., 2010; Jaudet & Hatey, 1984), however, there are exceptions. Indeed, previous studies in adult *D. tinctorius* demonstrate that, although corticosterone is the ACTH responsive glucocorticoid, adults release equal amounts of corticosterone and cortisol (Westrick et al. 2023), and increased cortisol in parents is associated with tadpole transport (Fischer & O’Connell, 2020). Taken together, these previous observations and our data here suggest that the production and relative abundance of corticosterone and cortisol may shift post-metamorphosis or as adults become sexually mature. Further exploration of these patterns and their potential functional consequences is an area of ongoing research.

### Conclusions

We found no significant differences in glucocorticoid excretion rates, whole body concentrations, or physiological responses to ACTH injection across tadpole developmental stages in *D. tinctorius*. Taken together, these results demonstrate an early-developed glucocorticoid stress response and suggest that the HPA axis is functional early in poison frog tadpole ontogeny. Our findings provide a foundation for exploring how *D. tinctorius* tadpoles physiologically respond to a habitat transitions and environmental stressors, as well as prepare for and undergo metamorphosis. Additionally, this pattern contrasts with other tadpole species, providing inroads for exciting comparative work to further build our understanding of how life-history shapes adaptive differences in the developmental timing of stress responses.

## Acknowledgements

We thank Drs. Britt Carlson and Nathan Schroeder from the Phenotypic Plasticity Research Experience for Community College Students (PRECS) program for adding Lauren Mobo to the team. We also thank Drs. Chris Holmes and Ken Paige, program directors of the Access and Achievement Program for adding Lucas Jimenez to the project. We appreciate the members of the Fischer Lab for feedback on previous versions of the manuscript and help with tadpole husbandry. Our work would not have been possible without the animal care support of the Division of Animal Resources (DAR) at UIUC.

## Author contributions

**Lisa L. Surber-Cunningham:** Conceptualization, Methodology, Formal Analysis, Investigation, Data Curation, Writing – Original Draft, Writing – Review & Editing, Visualization, Supervision, Project Administration. **Lucas S. Jimenez**: Investigation. **Lauren W. Mobo**: Investigation. **Sarah E. Westrick**: Conceptualization, Methodology, Formal Analysis, Writing – Review & Editing. **Eva K. Fischer**: Conceptualization, Methodology, Formal Analysis, Funding Acquisition, Resources, Supervision, Writing – Review & Editing.

